# Physiologically Relevant 3D CRISPR Screening Enhances Mechanistic Insight into Chemical Toxicity Compared to 2D Screening

**DOI:** 10.64898/2025.12.16.694776

**Authors:** Chanhee Kim, Zhaohan Zhu, Abderrahmane Tagmount, W. Brad Barbazuk, Rhonda Bacher, Christopher D. Vulpe

## Abstract

Many omics-based approaches in toxicology research primarily rely on correlative data, often lacking functional relationships or causal links between genotypes and phenotypes. CRISPR-based approaches can overcome this limitation by establishing direct causal connections between genes and toxicological phenotypes. Moreover, CRISPR screens enable scalable and systematic interrogation of gene function and associated mechanisms following chemical exposure, predominantly using *in vitro* models. In line with the paradigm of new approach methodologies (NAMs) in toxicology research, CRISPR screens hold promise to provide an *in vitro* cell-based functional toxicogenomics approach. One of the main limitations of conventional *in vitro* assays is their compromised physiological relevance to humans due to their inability to fully recapitulate *in vivo* physiology. To improve the functional and physiological relevance of the toxicogenomics approach, we developed a 3D CRISPR screening system using HepG2/C3A spheroids generated and cultivated in a continuously rotating bioreactor (ClinoStar). We first performed time-course 3D CRISPR screens to identify genes that confer growth disadvantage or advantage, influencing spheroid development compared to 2D cultures. We then applied this approach to a chemical toxicity study using doxorubicin, comparing the performance of the 3D and 2D systems in identifying chemical-specific mechanisms. The results showed that the 3D system captured more candidate genetic determinants and biological pathways related to DNA damage processes—a known toxicity mechanism of doxorubicin—demonstrating improved performance in identifying chemical-specific pathways over the 2D counterpart. In our screens, we employed custom CRISPR sgRNA libraries representing common human loss-of-function genetic variants (mean allele frequency > 0.1% in all individuals catalogued in the genome aggregation database), which potentially affect toxicity responses. By comparing our CRISPR screen results with previously reported genetic associations for doxorubicin response, we found that the 3D system identified more known associated genes than the 2D system. Together, the 3D CRISPR screening system demonstrated its feasibility and utility for physiologically relevant functional toxicogenomics. This platform enables *in vitro* NAMs, by providing a scalable and effective approach to identify causal genetic determinants and biological pathways that modulate chemical-induced toxicity.

## 1. Introduction

Systems biology approaches, such as multiple omics platforms, are widely applied in toxicological research to aid in the mechanistic understanding of chemical toxicity(Alexander-Dann et al., 2018; Harrill et al., 2021; Beale et al., 2022). However, such approaches primarily rely on correlative data, often lacking functional relationships or causal links between genotypes and phenotypes(Hasin et al., 2017) (e.g., correlation of gene expression and toxicity). Since the discovery of CRISPR(Jinek et al., 2012), this genome engineering tool is being widely adopted to interrogate gene function and its causal relationship to phenotypes across biomedical research(Barrangou and Doudna, 2016; Asmamaw and Zawdie, 2021); however, large-scale CRISPR screening applications in toxicology remain limited(Sobh and Vulpe, 2019; Sobh, Loguinov, Stornetta, et al., 2019; Sobh et al., 2021). CRISPR-based functional genomics screening (CRISPR screening) has transformed our ability to systematically interrogate gene function(Arroyo et al., 2016; Chang et al., 2020; Rossiter et al., 2021; Awah et al., 2022). By enabling genome-wide or targeted genetic perturbations with high efficiency, CRISPR screening provides a powerful and versatile platform to identify genetic determinants of diverse cellular processes at scale. In toxicology, CRISPR screens offer an unbiased approach to uncover the molecular mechanisms underlying chemical-induced toxicity and relevant adverse outcomes(Gaytán and Vulpe, 2014; Sobh, Loguinov, Stornetta, et al., 2019; Sobh, Loguinov, Yazici, et al., 2019; Wang et al., 2022). However, most CRISPR-based toxicity studies rely on 2D cell culture systems(Doench, 2018; Bock et al., 2022), which limit physiological relevance to human biology, often due to considerably different metabolic capacities as compared to *in vivo* systems(Duval et al., 2017; Kapałczyńska et al., 2018).

3D cell models offer significant advantages over conventional 2D cultures in toxicological research, as they better recapitulate human physiological conditions and toxicant responses, including improved xenobiotic metabolism that is more similar to *in vivo* metabolism(Gunness et al., 2013; Juarez-Moreno et al., 2022; Tang, Shi, et al., 2023). In addition, 3D cultures more closely mimic the *in vivo* microenvironment than 2D cultures through enhanced cell-cell and cell-extracellular matrix (ECM) interactions, tissue-like architecture, and gene and protein expression patterns that resemble *in vivo* states(Gunness et al., 2013; Kapałczyńska et al., 2018; Tang, Shi, et al., 2023). These *in vivo*-comparable characteristics, particularly those related to xenobiotic metabolism, enable 3D models to produce more physiologically relevant cellular responses than their 2D counterparts when the same cell lines are used(Kim et al., 2024, Anon, n.d.). Thus, 3D models may improve the identification of organism relevant responses critical for effective toxicological risk assessment. However, identifying functional gene–toxicant interactions in physiologically relevant human models remains challenging due to the lack of appropriate experimental systems. The recent shift toward reducing animal testing under the framework of new approach methodologies (NAMs) further underscores the importance of 3D *in vitro* models in toxicology(Stresser et al., 2024; Beilmann et al., 2025). While several applications of 3D CRISPR screening have been demonstrated in cancer research(Han et al., 2020; Takahashi et al., 2020), such approaches have not yet been established for toxicological studies.

Here, we developed a 3D CRISPR screening system for toxicological applications to enhance the physiological relevance of the conventional 2D functional toxicogenomics approach using HepG2/C3A human liver spheroids. First, we performed time-course 3D CRISPR screens to identify functional genetic components influencing spheroid growth compared to 2D culture in normal growth media. Next, we applied this approach to a chemical toxicity study using doxorubicin (Doxo) as a model chemical to compare the performance of 3D and 2D screens in identifying genes and pathways associated with Doxo-induced toxicity. Doxo was chosen due to its well-characterized DNA damage–inducing mechanism of action(Thorn et al., 2011; Nicoletto and Ofner, 2022), providing a reference framework for mechanistic comparison. We used custom CRISPR sgRNA libraries (PopVarLoF) representing common human loss-of-function (LoF) genes (mean allele frequency > 0.1% in all individuals catalogued in the genome aggregation database(Karczewski et al., 2020)) that could potentially affect toxicity responses. This screening approach was used to identify both known and previously unrecognized genes harboring common LoF genetic variants potentially modulating doxorubicin exposure-related phenotypes in people. Overall, we demonstrated that a 3D CRISPR screening system can provide a physiologically relevant and scalable platform for functional toxicogenomics, enabling the discovery of causal genetic determinants and pathways and complementing traditional genetic association studies.

## 2. Materials and methods

### 2.1 HepG2/C3A cell culture in 2D monolayer and 3D spheroid

The HepG2/C3A cells were directly purchased from the American Type Culture Collection (ATCC, Manassas, Virginia). Cells were maintained and passaged following ATCC’s recommended protocol. Minimum Essential Medium (MEM, Gibco) was supplemented with 10 % fetal bovine serum (FBS, Thermo Fisher Scientific, Waltham, MA), 1X MEM Non-Essential Amino Acids Solution 100X (Gibco, added for HepG2/C3A) and 1X Antibiotic-Antimycotic 100X (Gibco). Cells were cultured in a humidified incubator (Forma™ Series II Water-Jacketed CO2 Incubator, Thermo Scientific™) with 5% CO_2_ at 37 °C. For 3D spheroid cultures, we followed the protocol from our previous study(Kim et al., 2024) except that dissociation and reassembly of spheroids were added to the 3D CRISPR screening procedure which is described in 2.3. ClinoReactors and ClinoStar (CelVivo) were used to generate and maintain HepG2/C3A spheroids under a continuously rotating bioreactor system for up to 30 days.

### 2.2 Cytotoxicity of doxorubicin in 2D monolayer and 3D spheroid

The cytotoxicity of 2D HepG2/C3A cells exposed to Doxo was assessed using a range of nominal concentrations (0.4 - 100 µM) for 72 h by measuring ATP levels using the CellTiter-Glo2.0 cell viability assay kit (Promega, Madison, WI) following the manufacturer’s instruction. For 3D spheroid cytotoxicity assays, 10 spheroids on Day 7 were assessed using a range of nominal concentrations (0.4 - 100 µM) for 72 h by measuring ATP levels using the CellTiter-Glo 3D cell viability assay kit (Promega, Madison, WI) following the manufacturer’s instructions. The luminescence signals were read on a Synergy H1 microplate reader (BioTek Instruments, Winooski, VT). Relative luminescence signals were calculated and converted to % of viable cells compared to controls. The % values were used to determine the 25 % inhibitory concentrations (IC_25_) for Doxo in 2D and 3D using GraphPad Prism (version 10.1.0) with the dose-response sigmoid function.

### 2.3 CRISPR screens

Lentiviruses of the two custom PopVarLoF sgRNA plasmid libraries (SET1 and SET2, *manuscript under review*) were produced as previously described(Sobh, Loguinov, Stornetta, et al., 2019; Sobh, Loguinov, Yazici, et al., 2019) (a list of sgRNA sequences is in Table S1). The resulting PopVar LoF lentiviral libraries were functionally tittered in HepG2/C3A cells to determine the amount of virus required to obtain a multiplicity of infection of 0.3 for each library. For large-scale transduction of these libraries for 2D and 3D CRISPR screens, HepG2/C3A cells were seeded in four 12-well culture plates (1×10^6^ cells/well) prior to lentiviral transduction. After 24 h, polybrene (Sigma) was added to each well to a final concentration of 8 µg/ml, along with 0.5 µl of each PopVar LoF lentiviral library (SET1 and SET2), followed by centrifugation at 33 ◦C at 1000xG for 2 h (transduction). After incubation at 37 ◦C for 30 h, the media containing the lentiviral transduction mixture was replaced with 1 mL of the growth media in each well. After 24 h of recovery, we selected transduced cells with puromycin (2 μg/mL) until non-transduced controls were eliminated, ensuring that only successfully transduced cells remained. After 3 days of recovery and expansion from puromycin selection, the cells were used either for 2D monolayer screens or 3D spheroid screens. To generate transduced 3D spheroids, we cultured the transduced HepG2/C3A cells using the 3D culture method described in Section 2.1. Notably, the number of transduced cells corresponding to approximately 500-fold (500X) the total PopVar LoF library size was maintained throughout both 2D and 3D CRISPR screens(Doench, 2018). Two types of CRISPR screens were conducted in this study, as described below:

#### 2.3.1 Time-course screens 3D vs. 2D

Both transduced initial 2D monolayer cells and 3D spheroids (2D_Day 0 and 3D_ Day 0) were cultured in a CO_2_ incubator (T75 flasks) and ClinoStar (ClinoReactors) continuously for up to 30 days. Time-course analysis samples were harvested on Day 20 and Day 30 (2D_Day 20, 2D_Day 30, 3D_ Day 20, and 3D_ Day 30). Culture media were changed every 3-4 days for both 2D and 3D systems.

#### 2.3.2 Chemical toxicity screens (doxorubicin) 3D vs. 2D

Aimed to perform comparable CRISPR screens of chemical toxicity in 2D and 3D, we optimized the chemical exposure scenario of 3D according to our previous study(Kim et al., 2024), which indicated a reduced cell proliferation rate of 3D spheroids in continuous culture and suggested an appropriate time window for 3D CRISPR screen (Day 3 to Day 10). As described in Fig. 1B, HepG2/C3A 3D spheroids exposed to Doxo were dissociated into single cells using Accutase (STEMCELL Technologies) on Day 6 (when exposure was suspended), reassembled into spheroids on Day 7, and then re-exposed to Doxo until Day 14. This adjustment to the 3D exposure regimen ensured comparability with the 2D Doxo exposure condition in terms of the number of cell doublings under chemical exposure (selection). Doxo-exposed samples were harvested on Day 14 (7 cell doublings) for both 2D and 3D, with corresponding no-exposure controls.

**Fig. 1.**
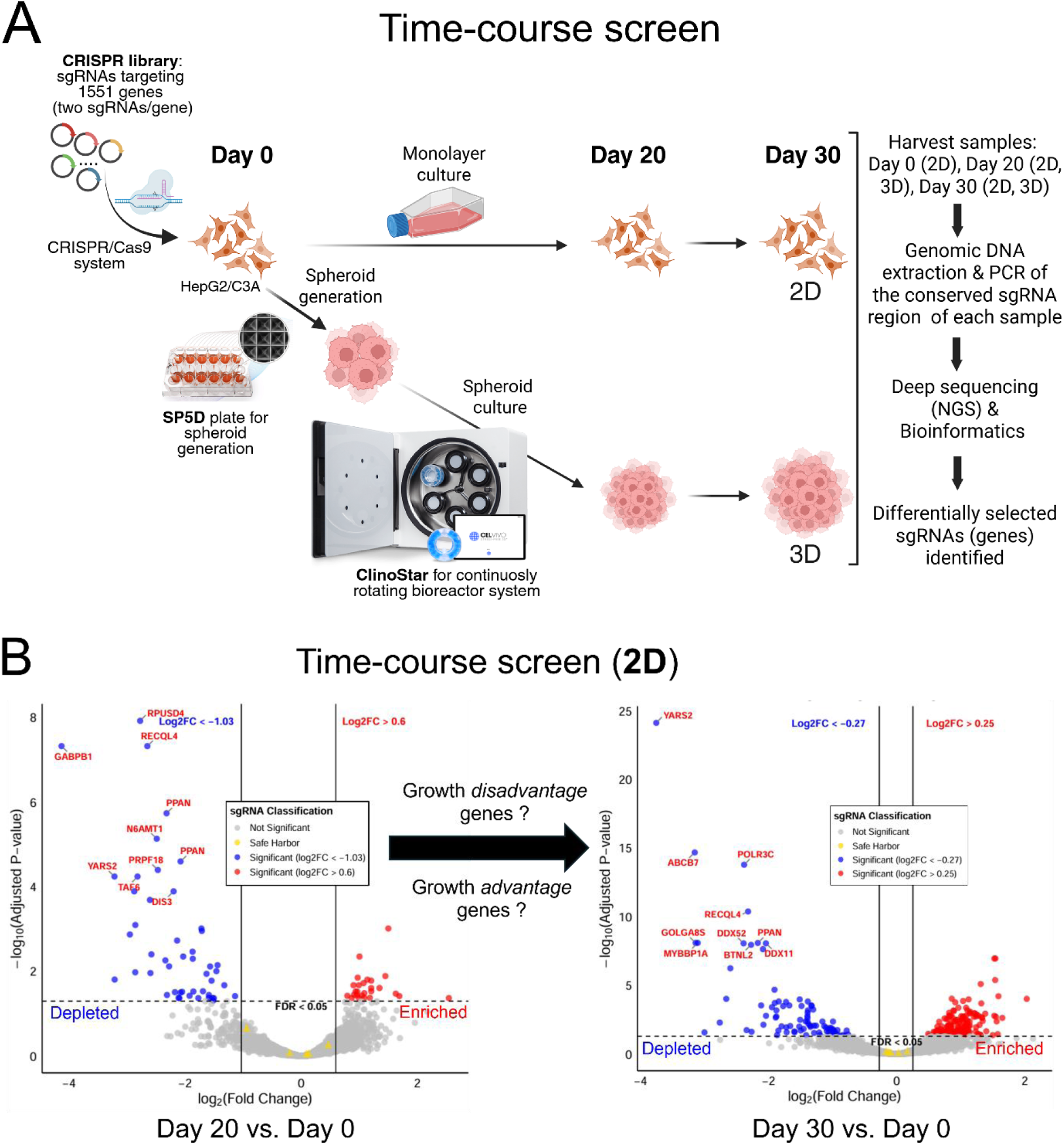
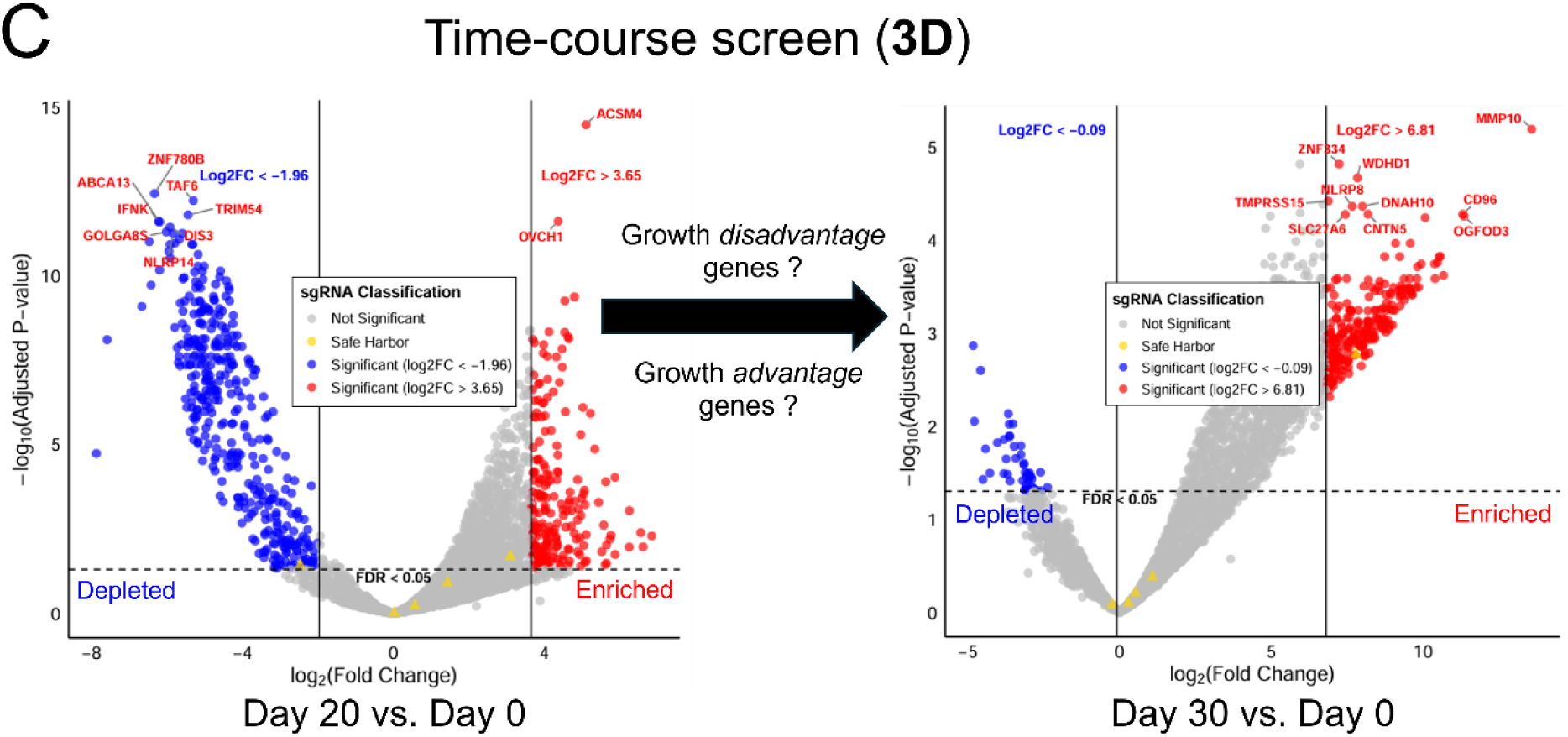
Time-course 3D CRISPR screening system compared to 2D screening in standard growth conditions. (A) Illustration of the workflow of 3D CRISPR screening system. SP5D, Spherical Plate 5D; NGS, next-generation sequencing. (B) and (C) Two volcano plots showing differentially selected candidate genes (depleted in blue; enriched in red) in 2D (B) and 3D (C) time-course CRISPR screens comparing Day 20 vs. Day 0 (left) and Day 30 vs. Day 0 (right). The top 10 statistically significant candidate genes (FDR<0.05) are labeled and sgRNA classification indicates not significant sgRNAs (grey) and safe harbor sgRNAs (yellow). By comparing two time-course volcano plots, sgRNAs (genes) that, when disrupted, resulted in a consistent and increasing growth disadvantage (depleted) or growth advantage (enriched) were identified. In each plot, x-axis indicates log_2_ (fold change) and y-axis indicates −log_10_ (adjusted p-value; FDR).

### 2.4 NGS preparation and sequencing

Genomic DNA was extracted from 1.6×10^6^ cells of each sample using the Quick-DNA Midiprep Plus Kit (ZYMO Research-D4075) following the manufacturer’s protocols. Amplicons for NGS Illumina sequencing were generated using pairs of universal CRISPR-FOR1 forward primer and CRISPR-REV**#** reverse primers (**#**: 1 to 48) each specific to a corresponding sample (Table S2). Amplicons for each sgRNA in each sample were then pooled and gel purified using the QIAquick Gel Extraction Kit (Qiagen) and quantified using the Qubit HS dsDNA assay (Thermo Scientific). Equimolar amounts of each amplicon library were multiplexed into a single pool, following the procedure described in previous studies(Sobh, Loguinov, Stornetta, et al., 2019; Sobh, Loguinov, Yazici, et al., 2019). The Illumina sequencing was carried out at the Interdisciplinary Center for Biotechnology Research (ICBR), University of Florida at Gainesville, using the NovaSeqX paired 150 bp high-throughput platform (Illumina).

### 2.5 Data analysis and bioinformatics

All FASTQ files were first assessed for sequencing quality using FastQC ( http://www.bioinformatics.babraham.ac.uk/projects/fastqc). Read counts were generated from FASTQ files using the MaGeCK count command(Li et al., 2014), which aligned reads to the provided PopVar LoF sgRNA library file and produced a sgRNA-level count matrix for downstream analysis. Differential selection analysis was performed using DESeq2 (version 1.40.2)(Love et al., 2014) in R (version 4.3.3) to identify differentially selected sgRNAs between the baseline (Day 0) and time-course points (Day 14 and Day 28). For each comparison of time-course screens (2D_Day 20 vs. Day 0, 2D_Day 30 vs. Day 0, 3D_ Day 20 vs. Day 0, and 3D_ Day 30 vs. Day 0), three Day 0 samples and two time-course samples from the respective time points were analyzed. For each comparison of chemical toxicity screens (2D_Doxo vs. 2D_Cont, 3D_Doxo vs. 3D_Cont), four 2D samples and two 3D samples were analyzed. The DESeq2 library size factor for normalization was calculated using the estimateSizeFactors function with the safe-harbor sgRNAs set as the control genes (Table S1). Safe-harbor control sgRNAs have been demonstrated to be a more appropriate baseline for normalization(Morgens et al., 2017; Chen et al., 2018). Differential selection was assessed using DESeq2’s negative binomial generalized linear model with the Safe-harbor-based library sizes included as an offset. We calculated the cumulative distribution function (CDF) of the effect sizes of negative controls (e.g., non-targeting sgRNAs and safe harbor controls) after normalization. The CDF was derived by first calculating the kernel density estimate (KDE) computed using the density function from the stats package in R and then converting the density into a cumulative distribution. We defined the 2.5% and 97.5% quantiles of this CDF as empirical cutoffs to identify significantly depleted or enriched effects. Adjusted p-values were calculated using the Benjamini–Hochberg procedure, and sgRNAs with an adjusted p-value < 0.05 were considered significantly differentially selected. For time-course screens, decreased sgRNA abundance indicated a growth-disadvantage phenotype, whereas increased abundance indicated a growth-advantage phenotype. For chemical toxicity screens, decreased sgRNA abundance indicated a sensitivity to Doxo, whereas increased abundance indicated a resistance to Doxo. Since the PopVarLoF library is composed of two separate libraries (SET1 and SET2) and each library has one sgRNA per gene (i.e., two sgRNAs per gene), a gene was deemed significant if at least one of its two sgRNAs was significantly differentially selected.

### 2.6 Functional enrichment analysis

Gene Ontology-Biological Process (GO-BP) and Kyoto Encyclopedia of Genes and Genomes (KEGG) pathway enrichment analyses were performed using the log2 fold-change (Log_2_FC) values of candidate genes, either sensitive or resistant to Doxo exposure in 2D and 3D screens. Enrichment analyses were conducted using the clusterProfiler tool implemented in SRPLOT software(Tang, Chen, et al., 2023) ([SRplot](https://bioinformatics.com.cn/en), accessed May 2025). STRING database(Szklarczyk et al., 2023) ([STRING: functional protein association networks](https://string-db.org/cgi/input?sessionId=bnf6XOFc9R0B&input_page_show_search=on), accessed Mar 2025) was used to cluster the gene products of candidate genes sensitive or resistant to Doxo exposure in 2D and 3D screens using a default setting (accessed on Mar 2025). We annotated each cluster containing two or more gene products based on GO-BP terms and KEGG pathways available in the STRING database.

### 2.7 ClinPGx doxorubicin phenotype data

The ClinPGx database ([ClinPGx](https://www.clinpgx.org/), accessed Sep. 2025) was queried to identify pharmacogenomic associations for anthracycline drugs, including doxorubicin, daunorubicin, epirubicin, and idarubicin, which share a common DNA damage–inducing mechanism of action. Curated datasets containing reported associations between genetic variants and drug-related phenotypes were downloaded from the database. To generate a non-redundant dataset, overlapping entries across the four anthracycline drugs were identified and removed before downstream comparisons.

### 3. Results

### 3.1. Time-course 3D CRISPR screen identified genes conferring growth disadvantage or advantage distinct from those identified in the 2D screen

To compare basal cellular physiology between the 3D CRISPR screening system and the conventional 2D system, we first performed time-course CRISPR screens under normal growth conditions to identify genes that, when disrupted, confer a growth disadvantage or advantage in each system (Fig. 1A). Using custom CRISPR sgRNA libraries representing the human genes with the most common aggregate LoF mutations (more detail in the introduction) developed in our previous study (*manuscript under review*), the initial mutant HepG2/C3A CRISPR library transduced and selected cell pool was used to establish either 3D spheroid cultures or 2D monolayer cultures (Day 0 for both), which were subsequently maintained for 30 days in their respective growth environments (Fig. 1A). For 3D culture, we employed the ClinoStar bioreactor system (CelVivo), which supports stable and continuous spheroid growth over extended periods, as demonstrated in our previous study(Kim et al., 2024). Time-course samples were harvested on Day 20 and Day 30 and compared with the Day 0 initial population. The relative abundance of each sgRNA, which is indicative of the abundance of each corresponding targeted mutant cell, was quantified using next-generation sequencing to determine how genetic disruption of each corresponding gene affected cell growth in both 3D and 2D conditions. Decreased sgRNA abundance indicates that disruption of the gene targeted by the sgRNA results in reduced growth or a growth-disadvantage phenotype. In contrast, increased abundance suggests that disruption of the gene targeted by the sgRNA results in enhanced growth, or a growth-advantage phenotype relative to other mutants in the pool. Differentially selected sgRNAs (genes) in 3D and 2D systems were identified across four comparison sets: Day 20 vs. Day 0 (3D), Day 30 vs. Day 0 (3D), Day 20 vs. Day 0 (2D), and Day 30 vs. Day 0 (2D) (Fig. 1B and Fig. 1C). Genes, when disrupted, showing increasing depletion over time, were empirically classified as growth-disadvantage genes, while those showing increasing enrichment were classified as growth-advantage genes in each respective system (Fig. 1B and Fig. 1C). By comparing relative sgRNA abundance (Log₂FC values) between time points and the Day 0 baseline, we identified 9 and 6 growth-disadvantage genes in 2D and 3D cultures, respectively, and 4 and 386 growth-advantage genes in 2D and 3D, respectively (Fig. 2A, Tables 1 and 2, Supplementary Table S3). For growth-disadvantage genes, we compared our results with the DepMap gene essentiality database(Tsherniak et al., 2017) (Table 1), which catalogs essential genes (EGs) critical for cell survival across diverse human cell lines. Among the 9 genes identified in 2D, 7 genes (*ABCB7*, *DDX11*, *DDX52*, *DIS3*, *MYBBP1A*, *POLR3C*, and *YARS2*) were previously reported as essential, one gene (*ADAM2*) was non-essential, and one gene (*GOLGA8S*) had no available data in DepMap (Table 1). For 3D growth-disadvantage genes, only 2 out of 6 genes (*CCDC63* and *TAF6*) were previously identified as essential, one gene (*FDXACB1*) was non-essential, and 3 genes (*MYH7B*, *NBPF9*, and *TLP1*) had no DepMap data (Table 1). Interestingly, while only 4 growth-advantage genes (*CYP2A13*, *KMT2C*, *OTOP3*, and *ZNF556*) were detected in the 2D screen, the 3D screen identified a substantially larger set of 355 growth-advantage genes (Table 2), allowing for meaningful functional enrichment analysis. STRING protein network analysis revealed that 3D growth-advantage genes were significantly enriched in the cytoplasm, cell periphery, membrane, and extracellular space (GO-CC terms; >50 genes per term; p < 0.05) (Fig. 2B), suggesting these genes are associated with cellular and extracellular compartments, and cell structure.

**Fig. 2.**
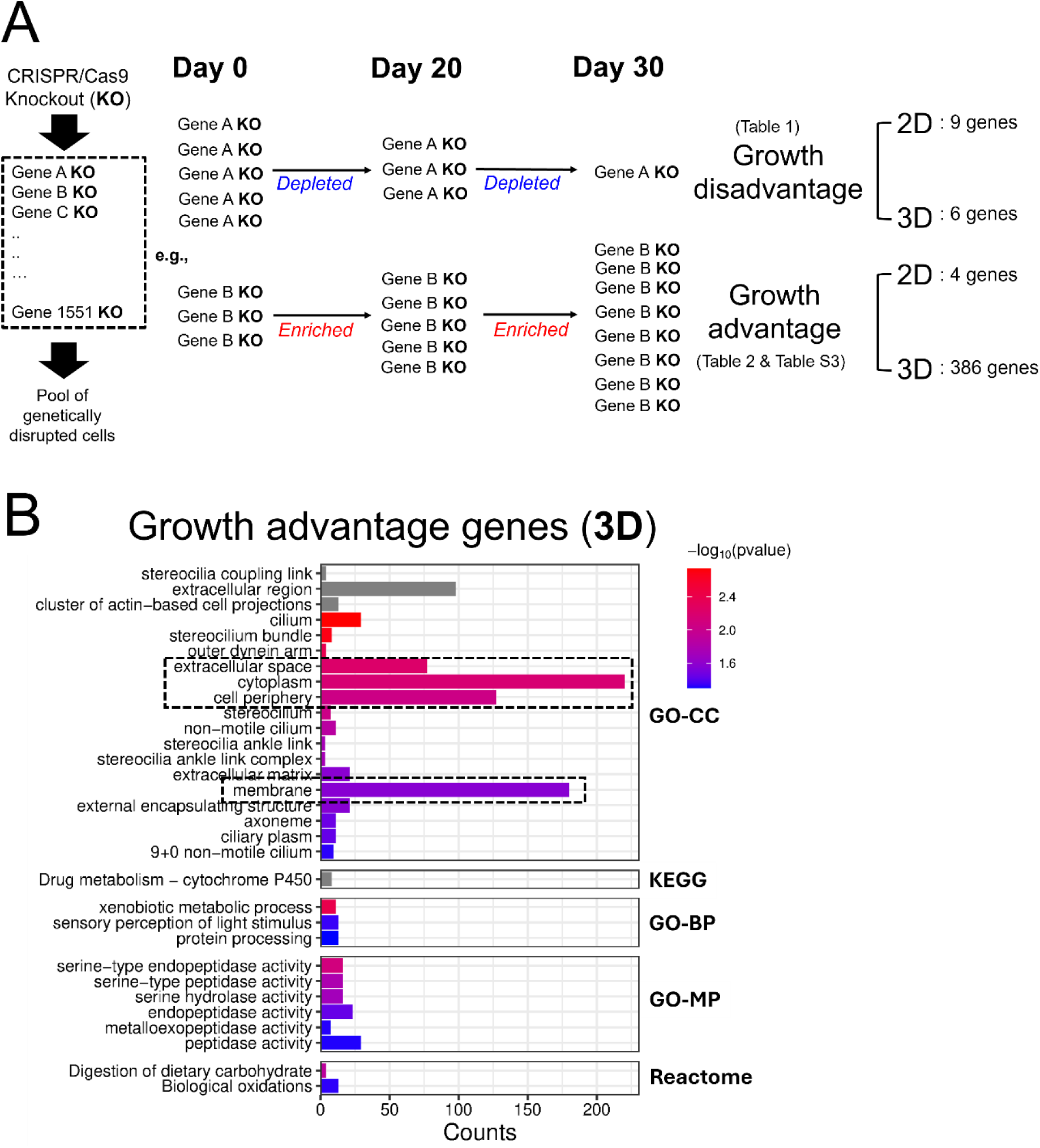
Differentially selected sgRNAs (genes) identified from time-course CRISPR screens of 2D monolayer and 3D spheroid in normal growth media. (A) Illustration of how growth disadvantage and advantage genes are defined. Both 2D and 3D CRISPR screens were conducted for 30 days. Time-course samples were collected on Day 20 and Day 30 and compared to the Day 0 initial mutant cell population. Gene KOs showing statistically significant and consistent depletion over time (based on Log_2_FC values for Day 30 vs. Day 0 and Day 20 vs. Day 0 comparisons) were classified as growth-disadvantage genes. In contrast, gene KOs showing continuous enrichment were classified as growth-advantage genes. (B) Functional enrichment of 3D growth-advantage genes (386 genes) using STRING protein network analysis. The dashed boxes indicate the terms with more than 50 gene counts and *p-value* less than 0.05. GO-CC, Gene Ontology-Cellular Component; GO-BP, Gene Ontology-Biological Process; GO-MP, Gene Ontology-Molecular Function.

**Table 1.** Comparison of growth disadvantage genes identified in time-course CRISPR screens in 2D monolayer and 3D spheroid systems with the gene essentiality profile catalogued in DepMap database(Tsherniak et al., 2017). Continuous depletion of a gene KO was empirically determined based on decreasing Log_2_FC values over time (Day 20 to Day 30) and gene KOs with such pattern were identified as growth disadvantage genes in the screens. Gene effect score less than 0 indicates gene essentiality where a score near −1 suggests that the gene is strongly essential (based on the Chronos dependency score from DepMap; [DepMap: The Cancer Dependency Map Project at Broad Institute](https://depmap.org/portal/))(Tsherniak et al., 2017).

|  | GENE | Day20_Log2FC<br>Vs. Day 0<br>(Empirical) | Day30_Log2FC<br>Vs. Day 0<br>(Empirical) | Growth<br>disadvantage<br>(Empirical) | Gene<br>Effect<br>(DepMap) | Gene<br>Essentiality<br>(DepMap) |
| --- | --- | --- | --- | --- | --- | --- |
| **2D** | ABCB7 | -2.86 | -3.15 | disadvantage | -1.90 | Essential |
|  | ADAM2 | -1.77 | -1.88 | disadvantage | 0.001 | **Not** essential |
|  | DDX11 | -1.32 | -2.04 | disadvantage | -0.67 | Essential |
|  | DDX52 | -1.51 | -2.39 | disadvantage | -0.47 | Essential |
|  | DIS3 | -1.87 | -2.09 | disadvantage | -0.64 | Essential |
|  | MYBBP1A | -2.95 | -3.11 | disadvantage | -0.46 | Essential |
|  | POLR3C | -1.71 | -2.38 | disadvantage | -1.14 | Essential |
|  | YARS2 | -3.22 | -3.75 | disadvantage | -1.02 | Essential |
|  | GOLGA8S | -2.58 | -3.14 | disadvantage | No data | No data |
| **3D** | CCDC63 | -2.91 | -3.15 | disadvantage | -0.21 | Essential |
|  | FDXACB1 | -3.17 | -3.51 | disadvantage | 0.01 | **Not** essential |
|  | MYH7B | -3.10 | -3.62 | disadvantage | No data | No data |
|  | NBPF9 | -3.43 | -3.73 | disadvantage | No data | No data |
|  | TAF6 | -2.74 | -3.80 | disadvantage | -1.65 | Essential |
|  | TLP1 | -2.18 | -3.25 | disadvantage | No data | No data |

**Table 2.** Comparison of growth advantage genes identified in time-course CRISPR screens in 2D monolayer and 3D spheroid systems. Continuous enrichment of a gene KO was empirically determined based on increasing Log_2_FC values over time (Day 20 to Day 30) and gene KOs with such pattern were identified as growth advantage genes in the screens. The entire list of 386 growth-advantage genes identified in the 3D system is available in Supplementary Table S3.

|  | GENE | Day20_Log2FC<br>Vs. Day 0<br>(Empirical) | Day30_Log2FC<br>Vs. Day 0<br>(Empirical) | Growth advantage<br>(Empirical) |
| --- | --- | --- | --- | --- |
| **2D** | CYP2A13 | 1.21 | 1.31 | advantage |
|  | KMT2C | 1.08 | 1.08 | advantage |
|  | OTOP3 | 1.21 | 1.53 | advantage |
|  | ZNF556 | 1.5 | 1.54 | advantage |
| **3D** | A2ML1 | 2.39 | 3.79 | advantage |
|  | ABCA10 | 1.99 | 4.40 | advantage |
|  | ABCB5 | 2.86 | 5.11 | advantage |
|  | ... | ... | ... | ... |
|  | UGT1A10 | 3.51 | 6.75 | advantage |
|  | UGT2A1 | 2.82 | 5.90 | advantage |

Together, our spheroid-based 3D CRISPR screening system, maintained in a continuously rotating bioreactor, not only supports extended spheroid culture but also identifies genes that specifically confer growth-disadvantage or growth-advantage phenotypes in 3D versus 2D environments under normal growth conditions. Notably, we observed a strikingly larger number of 3D-specific growth-advantage genes compared to 2D, highlighting unique effects of genetic disruptions on growth in spheroid cultures.

### 3.2. The 3D CRISPR screen identified more candidate genetic modulators of doxorubicin-induced toxicity than the 2D screen

To apply our 3D CRISPR screening system to the study of chemical toxicity mechanisms for comparison to the 2D system, we used doxorubicin (Doxo), which is a well-characterized DNA-damaging agent (Fig. 3A). IC_25_ concentrations of Doxo in 3D spheroid and 2D monolayer of HepG2/C3A cells were used to identify candidate genes that, when disrupted, increased sensitivity or resistance to Doxo-induced cellular toxicity (Supplementary Fig. S1). To ensure comparable chemical exposures across 3D and 2D systems the initial mutant cell populations were treated with Doxo for approximately seven cell doublings (14 days, collected on Day 14). For 3D cultures, HepG2/C3A spheroids were treated for 6 days, dissociated, and reassembled into new spheroids to match the effective doubling time of 2D cultures, thereby controlling for previously observed proliferation rate differences(Kim et al., 2024) (Fig. 3A, see Methods). Differentially selected sgRNAs (genes) were identified by quantifying relative sgRNA abundance via next-generation sequencing, revealing how genetic perturbations affected Doxo response in 3D versus 2D environments. Decreased sgRNA abundance indicated a sensitivity of the mutant cell with the corresponding gene disruption to Doxo, whereas increased abundance indicated a resistance of the mutant cell with the corresponding gene disruption to Doxo. The 2D Doxo CRISPR screen identified 60 candidate genetic modulators (41 sensitive genes and 19 resistant genes; Fig. 3B), whereas the 3D Doxo CRISPR screen identified 748 candidate modulators (91 sensitive genes and 657 resistant genes; Fig. 3C). Despite this large difference in the total number of modulators, 30 genes were consistently identified in both 2D and 3D screens, of which 28 genes conferred resistance to Doxo when disrupted (Fig. 3D). To identify enriched protein networks among the common genetic modulators, we performed STRING protein network analysis, which revealed a single enriched term, Ribosome Biogenesis, comprising four genes (*PPAN-P2RY11*, *NOC3L*, *DDX52*, and *PPAN*) shared between the 2D and 3D screens (Fig. 3D).

**Fig. 3.**
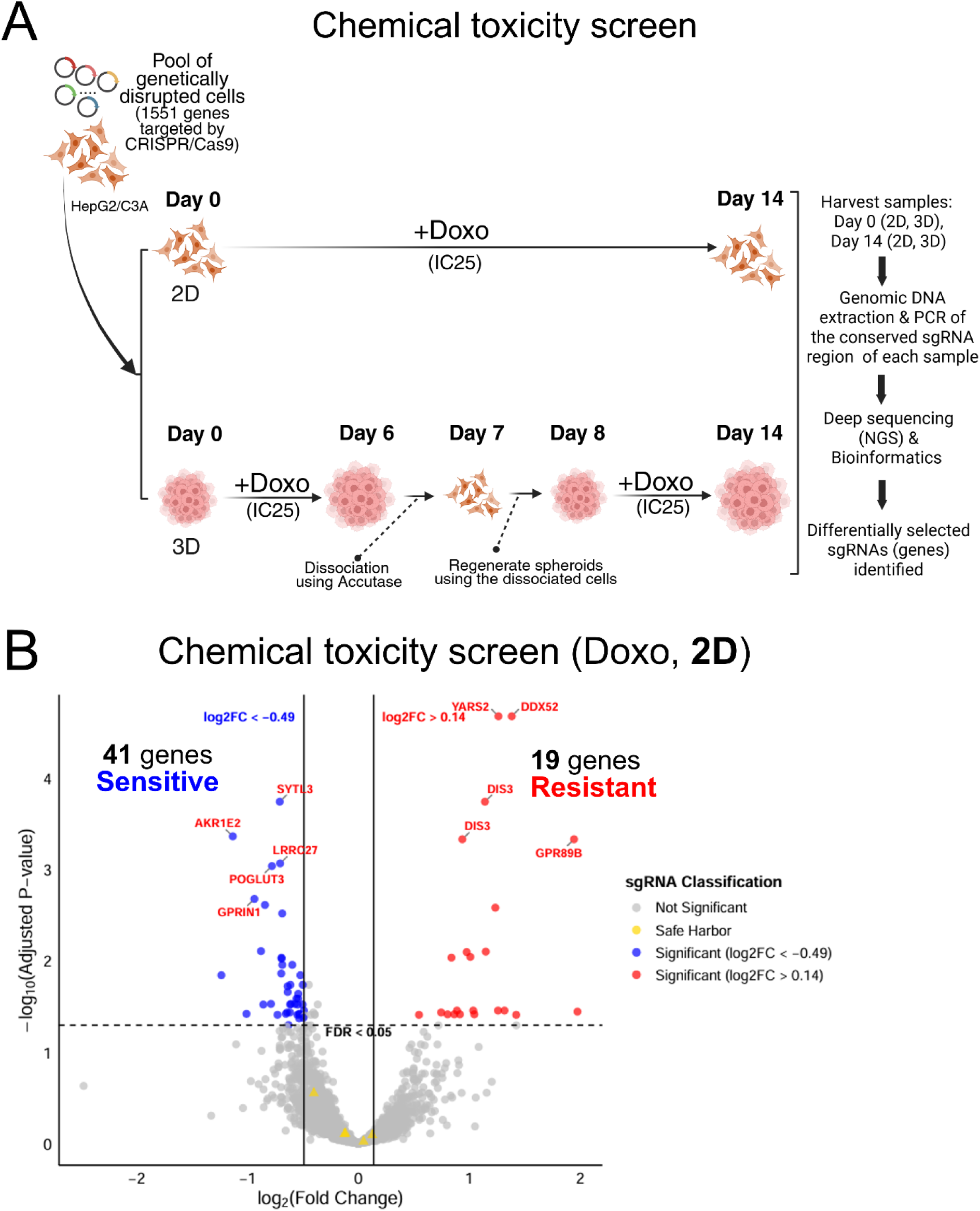
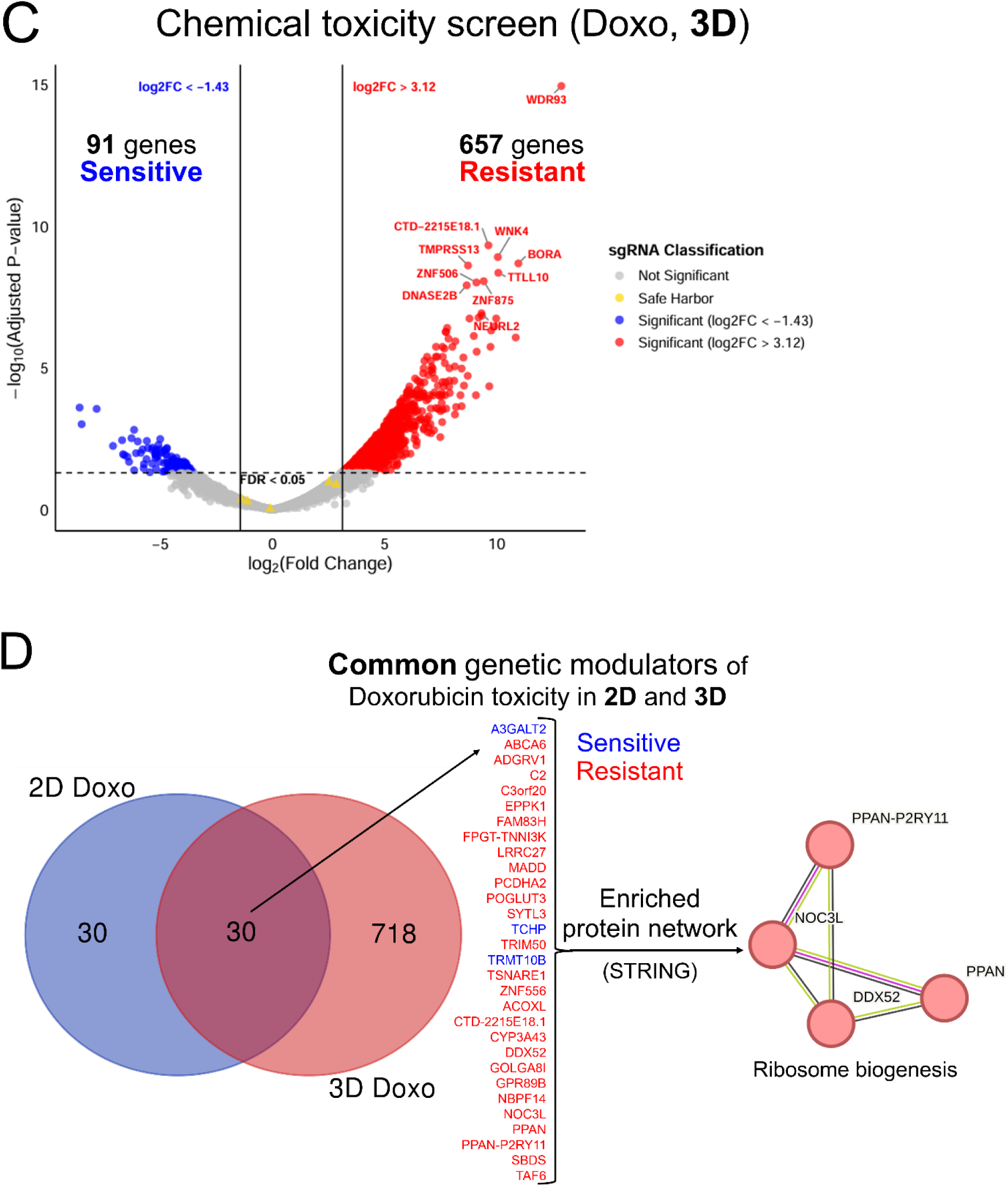
Application of the 3D CRISPR screening system to a chemical toxicity study using doxorubicin (Doxo) as a model chemical. (A) Schematic workflow of the 3D CRISPR screening system for Doxo-induced chemical toxicity screen compared to the 2D system. IC_25_, inhibitory concentration of 25, where a 25 % reduction in cell viability endpoints compared to the control is observed. (B) A volcano plot of 2D Doxo CRISPR screens identifies candidate genes that, when disrupted, result in increased Doxo sensitivity (blue dots) or increased Doxo resistance (red dots) in the CRISPR screens. (C) A volcano plot of 3D Doxo CRISPR screens displays Doxo-sensitive (blue dots) and Doxo-resistant (red dots) candidate genes identified in the screens. (B) and (C) The top 10 statistically significant candidate genes (FDR<0.05) are labeled and sgRNA classification indicates not significant sgRNAs (grey) and safe harbor sgRNAs (yellow). In each plot, x-axis indicates log_2_ (fold change) and y-axis indicates −log_10_ (adjusted p-value = FDR). (D) Common candidate genes identified in both 2D and 3D screens. A Venn diagram shows the number of common candidate genes. Blue and red colored genes indicate if disruption increases sensitivity or resistance to Doxo-induced cellular toxicity, respectively. An enriched functional protein network of the common candidate genes is visualized via STRING network analysis.

### 3.3. The 3D CRISPR screen of doxorubicin more accurately captured mechanisms relevant to chemical-specific toxicity compared to the 2D screen

In the 2D screens, Doxo-sensitive genes were enriched for cytoskeleton-related GO-BP terms, including intermediate filament cytoskeleton organization, intermediate filament-based processes, and apoptotic cell clearance (Fig. S2A). In contrast, 2D Doxo-resistant genes were primarily enriched for ribosome-related processes, such as rRNA processing, rRNA metabolic process, ribosome biogenesis, and ribosome assembly, as well as ncRNA-related processes including ncRNA metabolic process and ncRNA processing (Fig. S2B). KEGG pathway analysis revealed that 2D Doxo-sensitive genes were enriched for ABC transporters (hsa02010), Alcoholic liver disease (hsa04936), Glycosphingolipid biosynthesis (hsa00601), and Fatty acid biosynthesis (hsa00061) (Fig. 4A), whereas 2D Doxo-resistant genes were enriched for Nicotinate and nicotinamide metabolism (hsa00760), Basal transcription factors (hsa03022), and ABC transporters (hsa02010) (Fig. 4B).

**Fig. 4.**
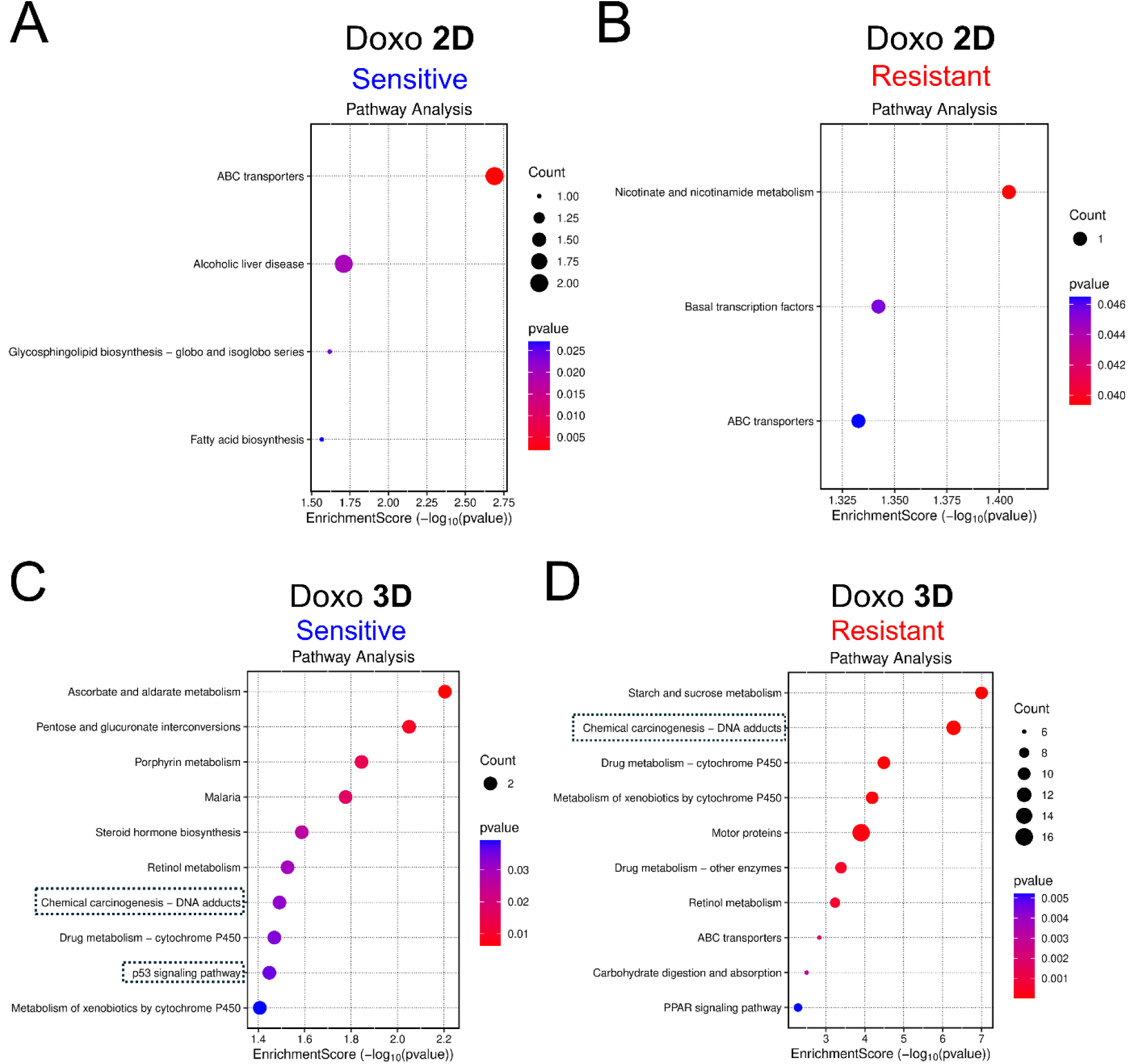
KEGG pathway enrichment of candidate genes modulating doxorubicin (Doxo)-induced cellular toxicity identified in 2D (A and B) and 3D (C and D) CRISPR screens. Results of the Doxo-sensitive genes are displayed with blue headers (A and C), while the resistant ones are displayed with red headers (B and D). Each plot presents the enrichment score on the x-axis while KEGG pathway terms (top 10 significant) are displayed along the y-axis. The size and color of the dots corresponding to the gene count and p-values, respectively. The dashed boxes indicate the terms directly related to DNA damage-response.

Given the substantially larger number of candidate modulators identified in the 3D screens, GO-BP and KEGG analyses captured more functional enrichment terms. For 3D Doxo-sensitive genes, the top GO-BP term was store-operated calcium entry (Fig. S2C). The 3D Doxo-resistant genes were enriched for processes related to general toxicant response and metabolism, including xenobiotic metabolic process, cellular response to xenobiotic stimulus, epoxygenase P450 pathway, long-chain fatty acid metabolic process, and fatty acid metabolic process (Fig. S2D). KEGG pathway analysis of 3D Doxo candidate genes revealed pathways directly related to DNA-damage response, a known Doxo-induced toxicity mechanism(Pfitzer et al., 2019) (Fig. 4C and 4D). 3D Doxo-sensitive genes were enriched for Ascorbate and aldarate metabolism (hsa00053) as the top term, and included Chemical carcinogenesis–DNA adducts (hsa05204) and p53 signaling pathway (hsa04115), indicative of Doxo-associated mechanisms(Cutts et al., 2005; Poirier, 2012; Yimit et al., 2019; Shen et al., 2023) (Fig. 4C). Similarly, 3D Doxo-resistant genes showed enrichment of Chemical carcinogenesis–DNA adducts (hsa05204), and additional pathways associated with general toxicant metabolism, including Drug metabolism–cytochrome P450 (hsa00982), Retinol metabolism (hsa00830), ABC transporters (hsa02010), and PPAR signaling pathway (hsa03320) (Fig. 4D). STRING protein network analysis of all 3D Doxo candidate genes identified several functional clusters, including glycogen metabolism, xenobiotic metabolism (Fig. S3A and Fig. S3B), and DNA damage response–related pathways (Fig. 5), which were not captured in 2D screens. Notably, the DNA damage response cluster included *RAD54L*, *GEN1*, *RECQL*, *POLN*, *FANCM*, and *FANCA* (Fig. 5A). STRING functional enrichment revealed that this cluster was significantly associated with multiple Reactome, KEGG, and GO-BP terms, including Fanconi Anemia pathway (Reactome and KEGG), DNA repair (Reactome and GO-BP), Double-strand break repair via homologous recombination (GO-BP), DNA-templated DNA replication (GO-BP), Interstrand cross-link repair (GO-BP), Double-strand break repair via synthesis-dependent strand annealing (GO-BP), Meiotic nuclear division (GO-BP), and Replication fork processing (GO-BP) (Fig. 5B).

**Fig. 5.**
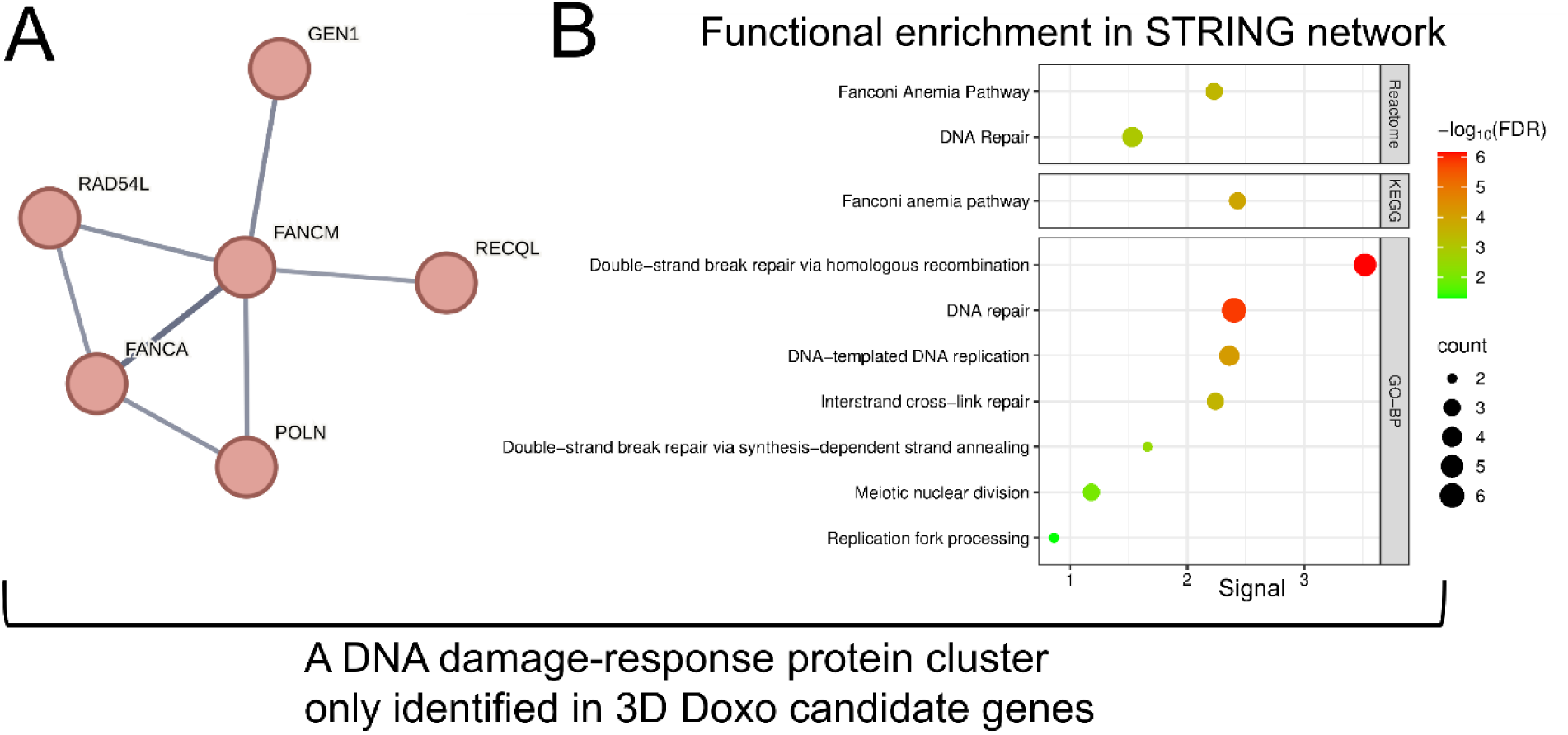
DNA damage response-related functional protein network significantly enriched in 3D doxorubicin (Doxo) candidate genes. (A) STRING protein network identified only in candidate genes of 3D Doxo CRISPR screens, not in the 2D screens. (B) Functional enrichment results reveal specific biological pathways related to DNA damage-response, including Reactome, KEGG, and GO-BP (Geno Ontology-Biological Process) terms. Each dot’s size and color corresponds to the gene count and FDR statistical significance, respectively.

Altogether, our 3D CRISPR screen not only identified a greater number of candidate genetic modulators of Doxo-induced toxicity (enhanced discovery potential) but also captured chemical-specific biological pathways, demonstrating enhanced mechanistic resolution compared to the 2D system.

### 3.4. Candidate modulators of doxorubicin toxicity in 3D screen showed greater overlap with known doxorubicin phenotypes than modulators from the 2D screen

In our CRISPR screens, we used a custom sgRNA library representing the genes with the most common aggregate loss-of-function (LoF) mutations in the human population as of GnomAD V3.0 (Mean aggregate allele frequency of >0.1%), which could influence toxicity response and susceptibility (*manuscript under review*). To evaluate the functional relevance of candidate modulators, we compared genes identified in the 3D and 2D Doxo CRISPR screens with previously reported clinical phenotypes associated with doxorubicin treatment. We queried ClinPGx, a comprehensive pharmacogenomics resource that catalogs curated associations between genes (genetic variants) and drug response phenotype(Gong et al., 2025), for doxorubicin as well as daunorubicin, epirubicin, and idarubicin, which share a common DNA damage–inducing mechanism, to capture a broad set of potential Doxo-related mechanistic phenotypes and their associated genetic variants.

Among 282 statistically significant gene–variant associations reported in ClinPGx, the 2D Doxo candidate genes matched 9 variants, all located in *CBR3* (Table S4, Fig. 6). In contrast, the 3D Doxo candidate genes matched 25 variants across 7 genes: *ABCC2* (10 variants), *ALDH3A1* (1 variant), *ATM* (2 variants), *CYP2C19* (6 variants), *GCKR* (2 variants), *GPR35* (1 variant), and *HMMR* (3 variants) (Table S4, Fig. 6). These results demonstrate that candidate modulators identified in the 3D CRISPR screen exhibit stronger concordance with established gene–Doxo phenotype associations than those identified in the 2D screen, highlighting the enhanced functional relevance and predictive power of the 3D system. We provide the LoF frequency data (Mean Allele Frequency from GnomAD V3.0) of candidate genes identified in our chemical screening, including the Doxo phenotype-associated genes (Table S5).

**Fig. 6.**
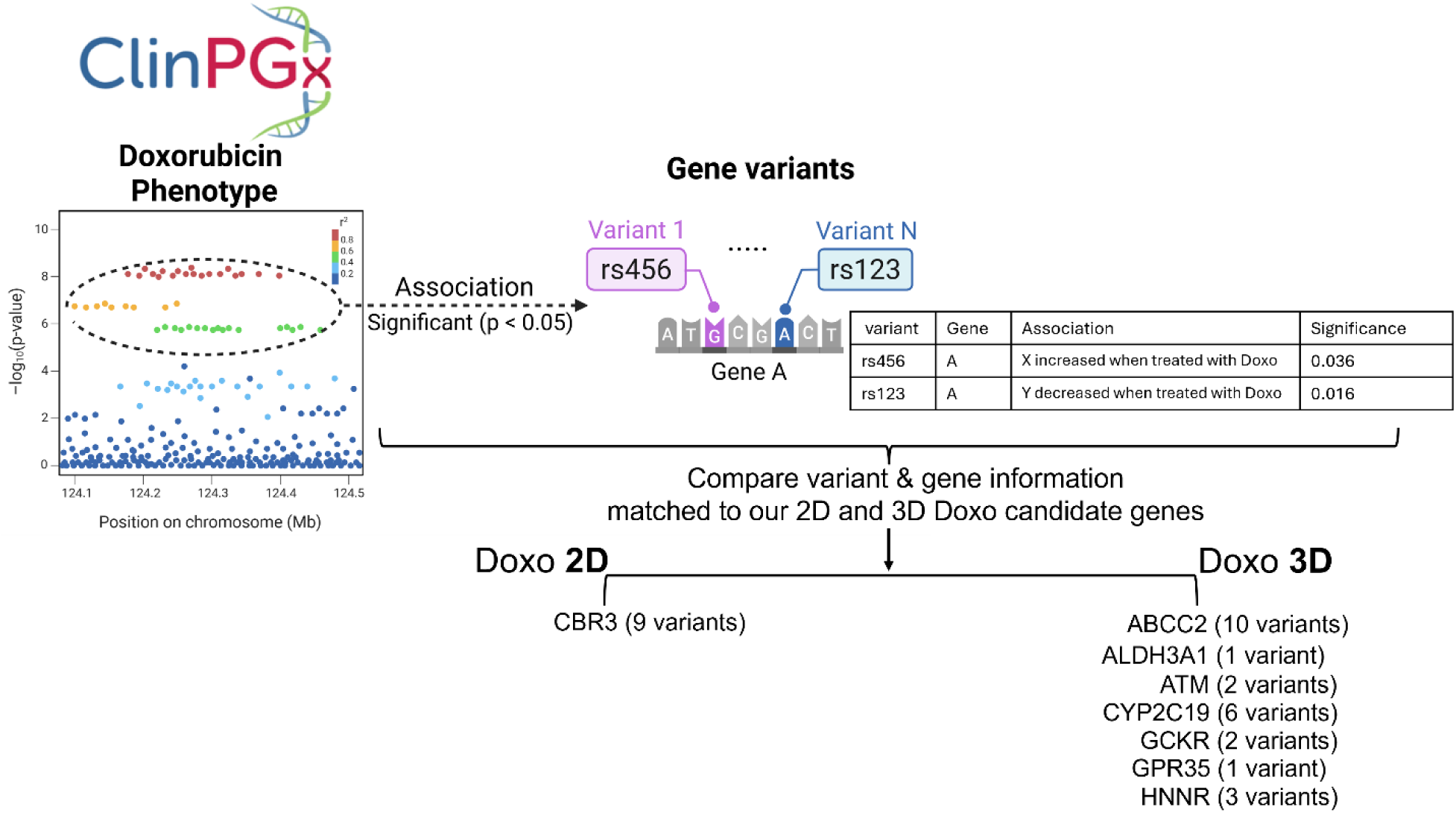
Comparison of candidate genetic modulators identified in 3D and 2D doxorubicin (Doxo) CRISPR screens with ClinPGx-reported gene–variant associations. Candidate genes identified in the 3D and 2D Doxo CRISPR screens were compared with genes and genetic variants previously associated with Doxo-related phenotypes using the ClinPGx database. The database was queried for four anthracycline drugs—doxorubicin, daunorubicin, epirubicin, and idarubicin—to capture a broad set of clinically relevant DNA damage–inducing phenotypes. Detailed information on individual variants, corresponding genes, associated phenotypes, and statistical significance values is provided in Table S4.

## Discussion

Here, we developed a 3D CRISPR-based genetic screening platform for functional toxicogenomics using an advanced 3D culture system. We applied and evaluated this platform in a chemical toxicity study using doxorubicin as a model toxicant. This work builds upon our recent study of 3D HepG2/C3A liver spheroids(Kim et al., 2024), in which we characterized the cellular and molecular features of a bioreactor-based spheroid culture system (ClinoStar). A critical exposure window with optimal cell proliferation was identified in the study, providing essential insights for optimizing the 3D CRISPR screening workflow for comparative chemical toxicity analyses(Kim et al., 2024). By combining bioreactor-based 3D spheroid culture with CRISPR-based functional genomics, our study advances NAM-based experimental approaches by providing a more physiologically relevant and genetically tractable platform to investigate chemical toxicity. The workflow performed in this study also enables systematic identification of functional genetic determinants of chemical responses within a more complex 3D culture systems, addressing key limitations of traditional 2D or single-endpoint NAM assays.

We observed substantial differences in both the number and types of candidate genetic determinants influencing normal growth and chemical toxicity responses between 3D spheroid and 2D monolayer systems. In the normal growth conditions, differentially selected sgRNAs (genes) in time-course CRISPR screens suggest unique and specific genetic components that are either disadvantageous or advantageous for cell growth in 3D and 2D. Notably, most genes conferring growth-disadvantage in 2D culture have been reported to be essential(Wang et al., 2015) in the DepMap database(Lagziel et al., 2019), but most 3D growth-disadvantage genes haven’t been documented as essential. This result not only supports the contention that our 2D time-course screen is comparable to previous 2D CRISPR screens in the existing genetic screening database in its gene essentiality data but also suggests that our 3D time-course screening could provide new and physiologically relevant gene essentiality information. The landscape of 3D gene essentiality has not yet been completely catalogued, partly due to technical and experimental limitations(Han et al., 2020; Grandhi et al., 2021). A growing body of evidence has suggested that 3D cell culture systems, compared to the 2D culture system, represent more accurately the microenvironment where cells reside in tissues(Edmondson et al., 2014; Law et al., 2021). This may explain difference of gene functional effects on cell growth in different microenvironments of the two systems in our screens. Context-dependent gene essentiality (i.e., growth disadvantage), where gene selection behavior varies depending on the growth medium and environment, has also been reported to arise from differences in cellular state(Lagziel et al., 2019; Rossiter et al., 2021) which could also support different cell behaviors between the 3D and 2D environment. Interestingly, we identified significantly more growth-advantage genes in 3D (355 genes) compared to 2D (4 genes). A STRING network analysis of these genes revealed functional enrichment in GO-CC terms associated with cell structure, likely reflecting spatial changes in cell-cell interactions(Alberts et al., 2002; Manz and Groves, 2010; Huang et al., 2022). Since structural components are required to maintain spheroid integrity, we interpret that these genes may be necessary for constraining cellular proliferation in 3D, although the exact mechanisms remain to be determined(Heinrich et al., 2020; Carpenter et al., 2023). A previous study(Han et al., 2020) focused on cancer dependency also reported that a CRISPR screen in 3D identified much more positive growth phenotypes (i.e., growth advantage genes) compared to the 2D screens, which is consistent with our findings. These findings also suggest that increased cell growth in spheroid systems of some mutants could represent an important caveat in the interpretation of 3D CRISPR screens.

To evaluate our 3D CRISPR screening platform relative to its 2D counterpart, we applied it to identify genetic modulators and functional biological pathways underlying chemical-induced toxicity using Doxo as a model toxicant. We chose Doxo due to its well-characterized DNA-damaging mechanism: it intercalates into DNA and inhibits topoisomerases, blocking replication, transcription, and translation, ultimately inducing apoptosis in rapidly dividing cells(Thorn et al., 2011, Anon, 2012). Despite its broad chemotherapeutic use, Doxo is associated with severe side effects, including cardiotoxicity(Wang et al., 2009; Al-Otaibi et al., 2022; Rodrigues et al., 2022; Robert Li et al., 2024; Stafford et al., 2024; Suga et al., 2025) closely linked to its interactions with iron metabolism(Ye et al., 2024) and although less widely appreciated, hepatotoxicity(Prasanna et al., 2020). For instance, clinical data indicate that serum aminotransferase levels, such as alanine aminotransferase (ALT) and aspartate aminotransferase (AST), are elevated in up to 40% of patients treated with Doxo, reflecting acute liver injury that is generally asymptomatic and transient(Anon, 2012). Hepatotoxicity remains a clinically relevant limitation(Anon, 2012; Prasanna et al., 2020; Lai et al., 2022) with its molecular mechanisms largely unexplored. Moreover, significant inter-individual variability in susceptibility to Doxo-induced hepatotoxicity and, cardiotoxicity suggests the presence of critical genetic determinants governing this adverse effect(Magdy et al., 2021; Xu et al., 2022; Fonoudi et al., 2024; El-Shorbagy et al., 2025a; Zobeydi et al., 2025).

In our screens, multiple novel candidate genes were identified exclusively in the 3D system, but not in conventional 2D cultures. Notably, we detected significantly more Doxo-resistant genes in 3D (657 genes) compared to 2D (19 genes), whereas the difference for Doxo-sensitive genes was smaller (91 vs. 41 genes). The reason for this pronounced disparity in resistant genes remains unclear; however, one possible explanation is that the 3D spheroid environment more accurately recapitulates cellular physiology, producing a stronger selective effect of Doxo on cell survival and proliferation over the course of spheroid growth and chemical exposure with the similar cytotoxic effect (i.e., IC_25_)(Fey and Wrzesinski, 2012; Juarez-Moreno et al., 2022). These findings also have potential relevance to the development of chemotherapeutic resistance and suggest that 2D CRISPR screens may not identify clinically relevant resistance mechanisms that could be important in the 3D growth environment of solid tumors.

In addition to the difference in numbers of candidate genetic modulators of Doxo-induced toxicity identified in our screens, functional enrichment analysis across culture conditions demonstrated that 3D CRISPR screens more accurately captured chemical-specific biological pathways. Several pathways directly related to DNA damage-response were enriched exclusively in 3D screens, including chemical carcinogenesis-DNA adducts and p53 signaling pathway. Furthermore, STRING protein network analysis of the 3D candidate genetic modulators of Doxo-induced toxicity revealed a cluster linked to DNA damage-response pathways, such as Fanconi anemia (FA) pathway, DNA repair, and homologous recombination (HR). FA is a rare genomic instability disorder, often caused by mutations in genes that regulate replication-dependent removal of DNA crosslinks(Jenkins et al., 2012). The FA pathway coordinates multiple DNA repair mechanisms, leading to the formation of repair structures in response to genotoxic insults, such as Doxo(Moldovan and D’Andrea, 2009). In our analysis, both *FANCA* and *FANCM* were identified within the functionally enriched protein cluster among 3D Doxo candidate genes; these proteins form part of the FA core complex(Moldovan and D’Andrea, 2009; Jenkins et al., 2012). The HR process is closely associated with *RAD51* expression, and *RAD54L*—a member of the DNA damage–response functional cluster identified here—actively facilitates HR(Alagpulinsa et al., 2014). Collectively, these findings support the contention that 3D CRISPR screens, at least for Doxo, may identify chemical-specific mechanisms of toxicity more accurately than 2D screens, establishing 3D screening as an effective functional toxicogenomics approach. The enhanced performance of 3D CRISPR screens observed here is consistent with previous studies in cancer biology(Han et al., 2020; Takahashi et al., 2020). For instance, Han et al. reported that CRISPR phenotypes in 3D cancer spheroids more accurately recapitulate *in vivo* phenotypes than 2D systems, highlighting the improved physiological relevance of 3D screens for capturing cancer-related phenotypes(Han et al., 2020). Similarly, Takahashi et al. showed that 3D CRISPR screens more effectively reveal the molecular pathogenesis mediated by nuclear factor erythroid 2-related factor 2 (NRF2) hyperactivation in lung cancer compared to their 2D counterparts.(Takahashi et al., 2020).

As we used custom CRISPR sgRNA libraries representing genes harboring the most common human genetic variants (PopVarLoF) in our screens, it was reasonable to compare our Doxo candidate genes from functional genetic screens with Doxo phenotypes reported in prior candidate gene-phenotype association or genome-wide association studies (GWAS). We acknowledge that our libraries do not cover the whole genome and thus might not capture all potential genes. We initially focused on the genes with the most common LoF mutations in the human population which thus could have population level relevance to clinical outcomes. We surveyed the ClinPGx database, which compiles genetic variants associated with Doxo treatment phenotypes, though most entries pertain to cardiotoxicity(WATANABE et al., 2023). Compared to the 2D candidate gene (one gene: *CBR3*), 3D candidate genes of Doxo-induced toxicity showed greater overlap with previously reported genes (7 genes: *ABCC2, ALDH3A1, ATM, CYP2C19, GCKR, GPR35,* and *HNNR*). Variants in genes involved in drug metabolism (*CBR3*(El-Shorbagy et al., 2025b)) and transport (*ABCC1*, *ABCB1*)(Magdy et al., 2022; El-Shorbagy et al., 2025b; Saito et al., 2025) have been associated with differences in drug response, treatment outcomes, and the risk of severe adverse effects. In particular, *CBR3* rs1056892(El-Shorbagy et al., 2025b) has been identified as major contributors to Doxo-induced cardiotoxicity, with emerging evidence suggesting that identification of such variants could guide the use of cardioprotective interventions(Magdy et al., 2021).

We also found additional candidates that appear relevant to Doxo phenotypes from a pathway-level perspective that had not been previously identified in association studies. For example, variations in genes encoding enzymes such as *UGT2B7* (involved in drug glucuronidation) are known to affect Doxo metabolism and clearance, thereby influencing hepatotoxic potential(Innocenti et al., 2001). Polymorphisms in *UGT2B7* have been shown to reduce glucuronidation activity, leading to increased systemic exposure to Doxo and a higher risk of liver injury(Hu et al., 2014, 2015). While *UGT2B7* itself did not emerge as a candidate genetic modulator in our screens, related genes, including *UGT2A1* (sensitive), *UGT2A2* (resistant), *UGT2B10* (sensitive), and *UGT2B11* (resistant), were significantly differentially selected (Table S4), which may reflect glucuronidation of different doxorubicin metabolites or potential changes in the extracellular matrix as suggested previously(Vitale et al., 2024). Notably, the entire set of 3D Doxo-sensitive genes was enriched for the pentose and glucuronate interconversions pathway (Fig. 4C), which is directly involved in glucuronidation(Kato et al., 2013; Oda et al., 2015). These findings suggest that follow-up investigations into functional or causal genetic variation-harboring genes identified in our screens (Table S4), which were preselected for population relevant genes with common LoF mutations, could improve the identification of predictive biomarkers for Doxo-induced liver toxicity that might be missed by conventional GWAS due to limited patient numbers, stringent statistical thresholds, or lack of functional validation(Wang et al., 2016; Schneider et al., 2017; Wells et al., 2017). Although not always systematic and scalable, functional validation has confirmed the biological relevance of several GWAS-nominated genes previously associated with Doxo-induced toxicity(Fonoudi et al., 2024). A notable example is a GWAS in 280 European-ancestry patients (32 cases vs. 248 controls) with childhood cancer, which identified a non-synonymous variant (rs2229774, S427L) in *RARG* as highly associated with Doxo-induced cardiotoxicity(Aminkeng et al., 2015). Gene-focused studies similarly highlighted the high interindividual variability in Doxo toxicity(Magdy et al., 2021), confirming the correlation of rs2229774 with adverse outcomes. Pre-chemotherapy screening for this variant, along with *RARG* agonist treatment, suggested potential cardioprotection in cancer patients. Furthermore, pathway enrichment analysis revealed that S427L cells exhibited significantly higher activation of Doxo-induced apoptosis, *TP53* targets, and oxidative phosphorylation compared to control cells, underscoring the mechanistic impact of this variant on drug response.

Currently available genetic approaches to address population variability and chemical susceptibility often lack diversity in study populations, limiting their ability to capture the full spectrum of variability. They also frequently face challenges in establishing causative relationships due to the inherent limitations of correlation-based methods(Amos et al., 2011). Our integrative CRISPR-based approach addresses these gaps by enabling prediction, prevention, and mechanistic understanding of Doxo-induced toxicity, as well as other liver-specific adverse outcomes or diseases. This approach bridges GWAS and gene-centric strategies by providing an intermediate, scalable method to identify population relevant functional genetic determinants that could influence response to toxicants, overcoming the limitations of each. Furthermore, several pharmacogenetic studies have highlighted the potential of genetic testing to risk-stratify breast cancer and lymphoma patients receiving doxorubicin-based chemotherapy(Jamieson and and Boddy, 2011; Bagdasaryan et al., 2022; Abdelfattah et al., 2025), emphasizing the importance of understanding genetic variants and their functional consequences, including chemical susceptibility. Collectively, these findings underscore the necessity of considering genetic variability when assessing hepatotoxicity risk in patients undergoing Doxo treatment. Promising experimental follow-up studies include single-cell level toxicogenomic profiling (single-cell RNA-seq) and high-content phenotypic assays (e.g., cell painting), both coupled with CRISPR library screening, to evaluate the chemical susceptibility derived from gene perturbations (i.e., genetic variability) at a single-cell resolution(Dixit et al., 2016; Bock et al., 2022; Feldman et al., 2022).

More broadly, the ability to systematically and scalably identify common genetic components—aggregate variants required for survival under distinct growth conditions, in the absence or presence of a toxicant—can enhance our understanding of functional genetic elements, context-specific susceptibility, regulatory networks, and toxicant-specific modulators across different culture systems.

## Conclusion

In this study, we demonstrate that a 3D spheroid-based CRISPR screening platform provides substantially improved mechanistic resolution for identifying genetic determinants of chemical-induced toxicity compared to conventional 2D monolayer screens. Time-course CRISPR screens under basal conditions revealed that 3D cultures operate under distinct physiological constraints, with markedly different patterns of growth-disadvantage and growth-advantage genes relative to 2D cultures. These differences highlight that cellular architecture, microenvironment, and metabolic state fundamentally alter genetic dependencies, underscoring the importance of physiologically relevant systems for functional genomics.

When applied to doxorubicin, a canonical DNA-damaging toxicant, the 3D CRISPR screen identified more than twelve-fold more candidate genetic modulators than the 2D screen and uniquely captured pathways consistent with established Doxo toxicity mechanisms—including Fanconi anemia signaling, double-strand break repair, replication fork processing pathways. In contrast, the 2D system predominantly recovered general stress-related processes rather than chemical-specific mechanisms. Importantly, candidate genes identified in 3D showed substantially greater overlap with known clinical pharmacogenomic associations for anthracycline toxicity, demonstrating improved alignment with human susceptibility and validating the translational relevance of the 3D platform.

Collectively, these findings show that 3D spheroid-based CRISPR screening markedly enhances the discovery of mechanistically meaningful gene–toxicant interactions and improves the detection of interindividual susceptibility factors compared to 2D approaches. Our results support the use of 3D genetic screening as a next-generation New Approach Methodology (NAM) for mechanistic toxicology and chemical safety evaluation. This platform offers a powerful strategy for defining causal genetic contributors to chemical responses, improving prediction of human-relevant toxicity pathways, and advancing the development of more accurate and physiologically grounded toxicological models.

## Supplementary material

Supplementary material is available at *Toxicology* online.

## Data availability

The raw and processed CRISPR screen genomics data with the corresponding metadata in this study will be deposited into Gene Expression Omnibus (GEO) database repository as soon as it is available.

