## Supplementary Figures for "Physiologically Relevant 3D CRISPR Screening Enhances Mechanistic Insight into Chemical Toxicity Compared to 2D Screening"

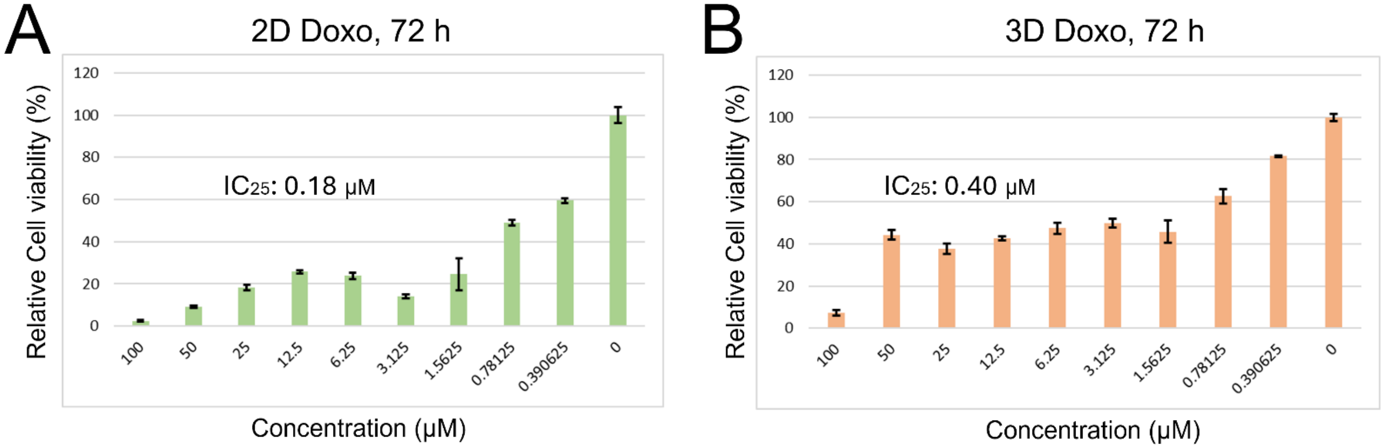


**Figure S1.** Cytotoxicity of doxorubicin (Doxo) in 2D and 3D cultures. Relative cell viability was measured after 72 h (3-day) exposure to increasing concentrations of Doxo (0–100 μM). $IC_{25}$ values for 2D and 3D cultures were derived using nonlinear regression analysis in Graphpad Prism software based on triplicate experiments.

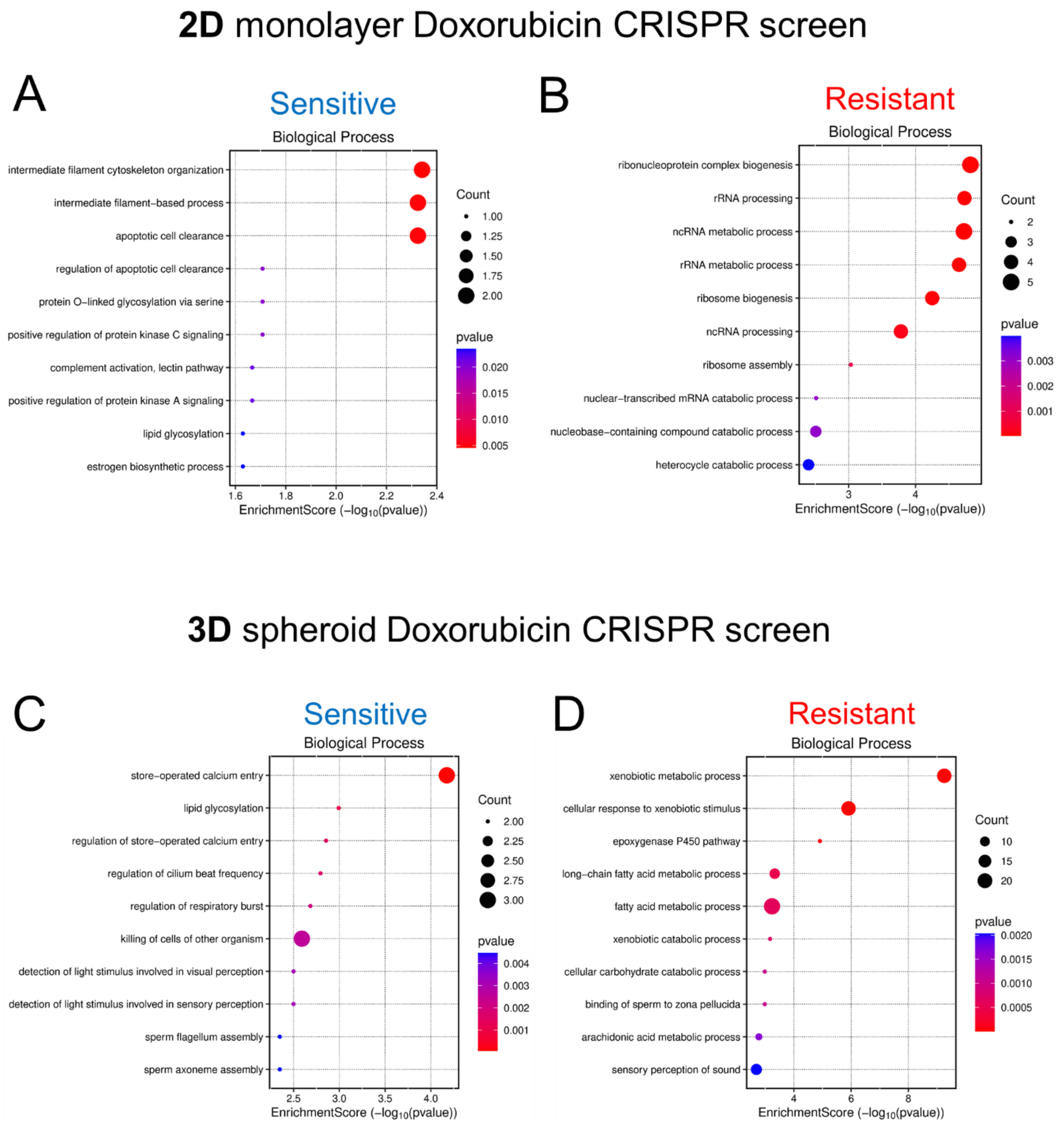


**Figure S2.** (A-D) Gene Ontology-Biological Process (GO-BP) enrichment analyses of candidate genes conferring Doxorubicin sensitivity or resistance identified in 2D and 3D CRISPR screens. The size of each bubble represents the number of candidate genes associated with a given GO-BP term (y-axis), while the x-axis indicates the enrichment score derived from $-\log_{10}$(p-value).

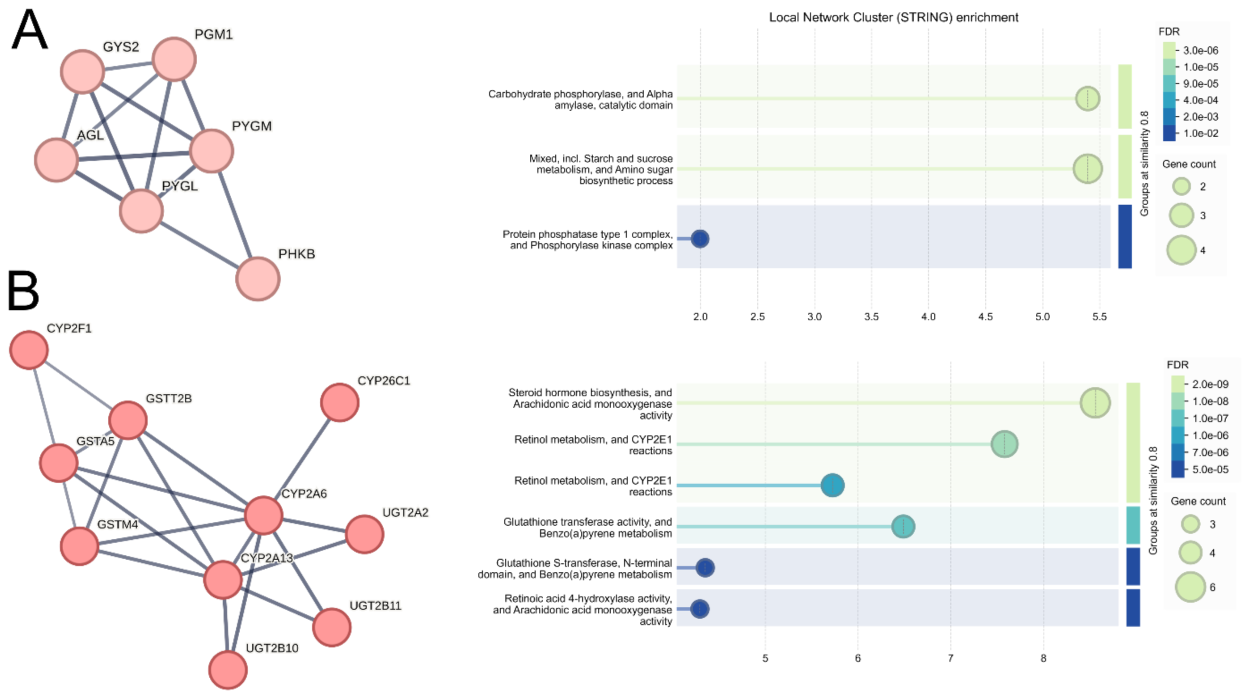


**Figure S3.** STRING network enrichment analysis of 3D doxorubicin (Doxo) candidate genes. Top 3 protein-protein interaction clusters identified from the 3D Doxo CRISPR screen are shown, in addition to the DNA damage-response cluster presented in Figure 5. (A) Glycogen metabolism-related network and (B) xenobiotic metabolism-related network are depicted with their corresponding enriched pathways.
